# Antibodies targeting conserved non-canonical antigens and endemic coronaviruses associate with favorable outcomes in severe COVID-19

**DOI:** 10.1101/2022.01.24.477545

**Authors:** Sai Preetham Peddireddy, Syed A. Rahman, Anthony R. Cillo, Godhev Manakkat Vijay, Ashwin Somasundaram, Creg J. Workman, William Bain, Bryan J. McVerry, Barbara Methe, Janet S. Lee, Prabir Ray, Anuradha Ray, Tullia C. Bruno, Dario A. A. Vignali, Georgios D. Kitsios, Alison Morris, Harinder Singh, Aniruddh Sarkar, Jishnu Das

## Abstract

While there have been extensive analyses characterizing cellular and humoral responses across the severity spectrum in COVID-19, predictors of outcomes within severe COVID-19 remain to be comprehensively elucidated. Recently, we identified divergent monocyte states as predictors of outcomes within severe COVID-19, but corresponding humoral profiles of risk have not been delineated. Furthermore, the nature of antibodies (Abs) directed against viral antigens beyond the spike protein or endemic coronavirus antigens and their associations with disease severity and outcomes remain poorly defined. We performed deep molecular profiling of Abs directed against a wide range of antigenic specificities in severe COVID-19 patients admitted to the ICU. The profiles consisted of canonical (S, RBD, N) and non-canonical (orf3a, orf8, nsp3, nps13 and M) antigenic specificities. Notably, multivariate machine learning (ML) models, generated using profiles of Abs directed against canonical or non-canonical antigens, were equally discriminative of recovery and mortality COVID-19 outcomes. In both ML models, survivors were associated with increased virus-specific IgA and IgG3 antibodies and with higher antigen-specific antibody galactosylation. Intriguingly, pre-pandemic healthy controls had cross-reactive Abs directed against nsp13 which is a conserved protein in other alpha and beta coronaviruses. Notably, higher levels of nsp13-specific IgA antibodies were associated with recovery in severe COVID-19. In keeping with these findings, a model built on Ab profiles for endemic coronavirus antigens was also predictive of COVID-19 outcome bifurcation, with higher levels of IgA and IgG3 antibodies against OC43 S and NL63 S being associated with survival. Our results suggest the importance of Abs targeting non-canonical SARS-CoV-2 antigens as well as those directed against endemic coronaviruses in favorable outcomes of severe COVID-19.

## Introduction

The continued spread of SARS-CoV-2 remains a significant threat globally, in spite of deployment of effective vaccines, due to newly emerging variants. A critical challenge is posed by the high symptomatic heterogeneity and unpredictable course of disease progression in COVID-19 (Rodebaugh et al., 2021).This contributes to the overburdening of health care systems leading to significant additional morbidity and mortality. Progression from asymptomatic infection or mild symptoms to severe disease has been broadly linked to advanced age and certain comorbidities (Ng et al., 2021). However, for those with severe COVID-19 disease, there is still a lack of personalized predictors of the course of disease and its outcomes. Although, significant effort has been dedicated to establishing the immunological underpinnings of COVID-19 (Carvalho et al., 2021) but the immunological drivers of mortality and survival outcomes within severe COVID-19 patients remain unclear. Recently, we identified dysregulated monocyte states as key predictors of outcomes within severe COVID-19 (Cillo et al., 2021). However, the corresponding antibody profiles that can predict COVID-19 mortality outcomes have yet to be delineated.

The humoral response directed against selected SARS-CoV-2 antigens, e.g., Spike (S) and Nucleocapsid (N), or their sub-domains, e.g., Receptor Binding Domain (RBD) of S, which taken together we term as canonical antigens here, have been extensively studied (Atyeo et al., 2020; Bartsch et al., 2021; Zohar et al., 2020). Relative to those with asymptomatic infection or with mild symptoms, antibody titers against canonical antigens are higher in patients with severe disease leading to early concerns about antibodies contributing to disease pathology, potentially via mechanisms like Antibody Dependent Enhancement (ADE) (Iwasaki and Yang, 2020; Lee et al., 2020) or via the antibody-mediated activation of inflammatory pathways especially since proinflammatory antibody Fc structures have been found to correlate with disease severity (Bye et al., 2021; Chakraborty et al., 2020; Hoepel et al., 2021; Larsen et al., 2021). Vaccine studies, meanwhile, have shown that titers of vaccine-elicited neutralizing antibodies directed against the S antigen are a key correlate of protection (Khoury et al., 2021; Sadarangani et al., 2021). Recently, longitudinal profiling of antibodies against canonical antigens, after natural infection, has revealed distinct temporal trajectories of immunoglobulin (Ig) subclasses and non-neutralizing functions of these antibodies that track with disease severity and outcome (Zohar et al., 2020). However, it remains to be determined which of these features of Abs directed against canonical antigens are predictive of recovery from severe COVID-19 disease.

Beyond the canonical antigens, the SARS-CoV-2 genome is predicted to encode up to 25 additional proteins (Gordon et al., 2020), which we term here as non-canonical antigens. It has been observed that cellular and humoral immune responses directed against these non-canonical targets also arise upon SARS-CoV-2 infection (Grifoni et al., 2020; Shrock et al., 2020). Antibody responses against some non-canonical antigens have been shown to be serological markers of COVID-19 at early and late time-points of illness (Hachim et al., 2020). However, it remains to be determined if antibodies directed against non-canonical antigens versus those directed against canonical antigens can independently or combinatorially predict the outcomes of severe COVID-19 disease. Given the prolonged exposure to a high viral burden in patients with severe COVID-19, antibodies against non-canonical antigens may play a role in protection or exacerbation in severe COVID-19. In the context of other viral infections, generation of antibodies directed against non-neutralizing targets has been linked to either protective, neutral or detrimental effects (Lamere et al., 2011; To et al., 2012). Indeed, in the context of COVID-19 as well, there is evidence pointing to the lack of selective targeting of S versus N antigens being linked to disease severity (Zohar et al., 2020). Additionally, it is notable that a number of these non-canonical SARS-CoV-2 protein antigens are known to share high sequence similarity with the corresponding proteins in endemic human coronaviruses (eHCoVs) (Hicks et al., 2021). Whether prior eHCoV exposure and the associated immune memory effects outcome after SARS-CoV-2 infection remains an unsettled debate and both protective and pathological effects have been reported in recent literature (Aydillo et al., 2021; Guo et al., 2021). It is plausible that the effects of prior eHCoV infection is mediated via the evolution or expansion of the pre-existing or cross-reactive B cell clones to these highly similar non-canonical antigens. Thus, tracking the antibody responses against these antigens could provide important insights in formulating improved SARS-CoV-2 as well as pan-coronavirus vaccines beyond the S-based formulations in current use.

To address these questions regarding the association and potential functions of specific Abs in severe COVID-19 disease outcomes, we developed a highly multiplexed, sample-sparing SARS-CoV-2 humoral profiling platform which measures biophysical properties of antigen-specific antibodies directed against a broad set of canonical and non-canonical antigens as well as eHCoV antigens including their isotypes, subclasses, Fc receptor binding and glycosylation. Serum samples of COVID-19 patients admitted into the ICU were profiled using this antibody profiling platform along with those from pre-pandemic healthy controls. The resultant high-dimensional data was analyzed with machine-learning based methods to generate multivariate models of Ab features that predicted severe COVID-19 disease outcomes. Importantly, we found that Abs directed against canonical and non-canonical antigens were independently equally predictive of disease outcomes. We also found that similar antibody profile differences of both antigen classes drove outcome bifurcation with survivors having more IgA and IgG3 antibodies and higher antigen-specific antibody galactosylation. Notably, pre-pandemic healthy controls were found to have antibodies against specific non-canonical antigens with high similarity to those in eHCoVs. Finally, eHCoV-specific antibodies were themselves also predictive of outcome in severe COVID-19 with higher levels of IgA and IgG3 being associated with survival. Thus, our results suggest the importance of Abs targeting non-canonical SARS-CoV-2 antigens as well as Abs directed against endemic coronaviruses in favorable outcomes of severe COVID-19.

## Results

### Multivariate antibody responses against canonical antigens are predictive of severe COVID-19 outcomes

Using our highly multiplexed, SARS-CoV-2 antibody profiling platform, we characterized and quantified serum Abs directed against canonical antigens for 21 severe COVID-19 patients from blood drawn soon after their ICU admission (Fig. 1a, UPMC cohort). Importantly, all 21 patients represented severe cases from natural infection (14 survivors and 7 non-survivors). Demographic details for these patients have been previously described (Bain et al., 2021; Cillo et al., 2021). Briefly, COVID-19 was diagnosed in these subjects based on reference-standard nasopharyngeal swab SARS-CoV-2 qPCR. Patients were admitted to the ICU a median of 6 days after symptom onset and serum was collected from these patients within 24 hours post-enrollment in the study. Of a wide range of measured clinical covariates, higher age and higher BMI showed trends of being associated with higher mortality and the administration of glucocorticoids trended to being associated with survival, none of these were univariate significant at these sample sizes (Bain et al., 2021; Cillo et al., 2021). This further motivates the need for the use of a multivariate approach that relies on molecular rather than clinical features to predict outcome bifurcation.

**Figure 1.**
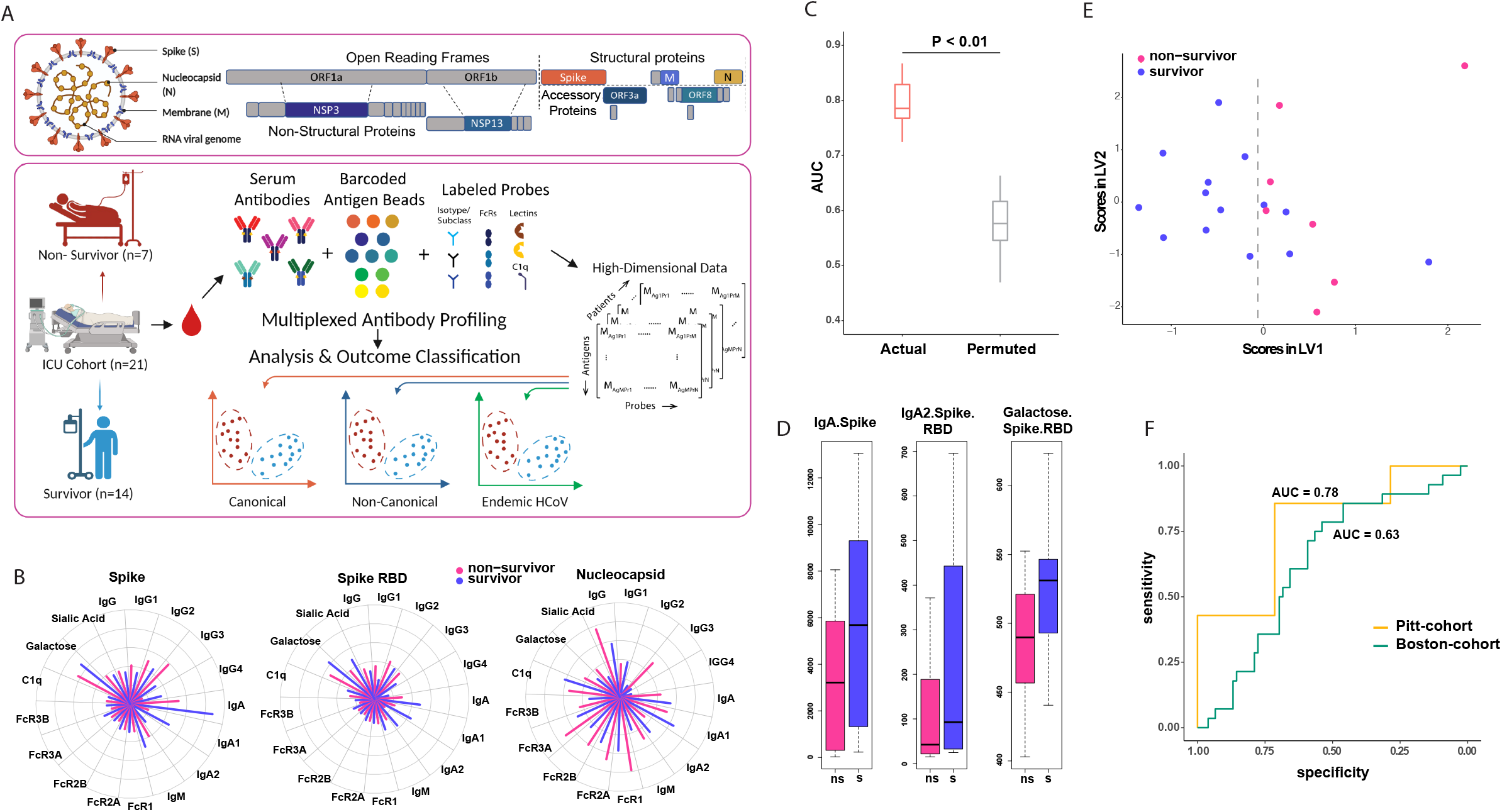
Multivariate antibody responses against canonical antigens are predictive of severe COVID-19 outcomes. A. Conceptual overview of SARS-CoV-2 antibody profiling platform characterized and quantified serum Abs directed against canonical/non-canonical antigens. B. Polar plots illustrating measured Ab features against canonical Ag specificities – Spike, Spike RBD, and Nucleocapsid C. Performance of LASSO model to discriminate between survivors and non-survivors built using deep humoral profiles against canonical Ag specificities. Model performance is measured in a k-fold cross-validation framework with permutation testing. Actual denotes the performance of the model, built on real data. Permuted denotes performance of the model on shuffled data in a matched cross-validation framework (negative control). D. LASSO-selected features from model built using deep humoral profiles against canonical Ag specificities. E. PLS-DA using only the LASSO-selected features from the model in (D) to discriminate between survivors and non-survivors. F. Performance of the model (built using the Pitt cohort) on an orthogonal (MGH, Boston) cohort.

Unlike well-characterized differences in Spike antibody titers of COVID-19 patients across the severity spectrum, we wished to determine whether antibody profiles at the point of ICU admission could predict bifurcation of subsequent outcomes – survival vs death (Fig. 1a). Furthermore, unlike previous studies that have focused on temporal differences (Zohar et al., 2020), we focused on Ab profiles measured at the point of ICU admission as it represents a clinically relevant and actionable time-point. Further, there weren’t significant differences in other characteristics including age, treatment and viral loads between the survivors and non-survivors.

We observed that both survivors and non-survivors had significantly higher antibodies across canonical specificities than pre-pandemic healthy controls, confirming the quality and specificity of our assay (Figs. S1a-S1o). However, the univariate differences between survivors and non-survivors were not striking (Fig. 1b, Figs. S1a-S1o). No single feature was discriminative by clinical outcomes. Therefore, we pursued a multivariate machine learning approach that incorporates different quantitative and qualitative aspects of the antibody response to determine if it could discriminate patients by clinical outcome. We used a two-step machine learning approach, as previously described (Ackerman et al., 2018; Das et al., 2020; Lu et al., 2020a; Sadanand et al., 2018; Suscovich et al., 2020), in an attempt to identify a minimal set of predictive Ab features that could discriminate between survivors and non-survivors. Our approach comprised feature selection using the LASSO; the use of L1 regularization on high-dimensional data (i.e., data where the number of Ab features far exceeds the number of subjects) that helps prevent overfitting (Ackerman et al., 2018; Das et al., 2020; Lu et al., 2020a; Sadanand et al., 2018; Suscovich et al., 2020). This was followed by classification using the down-selected features. We found that a model generated using this approach was significantly predictive of outcome as measured in a k-fold cross-validation framework with permutation testing (Fig. 1c, Methods). The model was based on three features – anti-Spike IgA, anti-Spike RBD IgA2 and RBD-directed antibody galactosylation (Fig. 1d). As an orthogonal way to visualize the stratification achieved by these Ab features, we performed partial least squares discriminant analyses (PLS-DA) using just these three down-selected features. The PLS-DA demonstrated that these 3 markers were able to stratify the survivors and the non-survivors (Fig. 1e). Notably, all three Ab features were higher in survivors compared to non-survivors (Fig. 1d), suggesting that higher levels of IgA antibodies directed against the Spike protein or its RBD and the increased levels of galactosylation of RBD-specific antibodies are associated with favorable outcomes.

To validate the robustness of the uncovered Ab features, we tested the performance of our model on an orthogonal cohort of severe ICU patients (MGH cohort) (Zohar et al., 2020). Critically, we had no role in the study design or recruitment strategy for this cohort. Our model generated using the UPMC cohort remained significantly predictive for the MGH cohort (Fig. 1f), albeit with a slightly decreased performance. We attributed this reduction in performance to the availability of fewer features in the MGH cohort dataset, specifically the lack of antigen-specific antibody glycosylation measurements, one of the three discriminating features for our model. The other 2 features (anti-Spike IgA, anti-RBD IgA2) exhibited identical univariate trends across the two cohorts and datasets (Fig. S1p). Overall, our results demonstrate that a model built using Ab profiles corresponding to canonical specificities is robust, both to cross-validation and cross-prediction with a distinct cohort. More importantly, they demonstrate that within severe COVID-19 patients, outcome bifurcation can be accurately predicted at the point of ICU admission based on IgA rather than IgG antibodies directed against S and its RBD as well as RBD-specific antibody galactosylation.

### Multivariate antibody responses against non-canonical antigens independently predict severe COVID-19 outcomes

Next, we sought to examine the levels of Ab responses directed against non-canonical antigens (nsp3, nsp13, orf3a, orf8) in patients with severe COVID-19 and whether they were independently predictive of outcome. We detected Ab responses to these non-canonical antigens both in the survivors and non-survivors, with no significant univariate differences between them (Fig 2a, Figs. S2a-S2o), similar to what we observed for canonical Ab specificities. So, we constructed a multivariate model, as described above, using only the Ab responses corresponding to non-canonical specificities. Strikingly, this model was also significantly predictive of outcomes (Fig. 2b), and the performance of this model was as good as that of our previous model built using canonical specificities (Figs. 2b, 1c). The model selected four features: anti-orf8 IgA, anti-nsp13 IgG3, anti-M antibody FcR3A binding and anti-M antibody galactosylation (Fig. 2c). Analogous to our earlier analyses, a PLS-DA visualization also demonstrated that these 4 features were able to stratify the survivors and the non-survivors (Fig. 2d). Importantly, these 4 Ab features, unlike many others (Figs. S2a-S2o), were higher in survivors. Our findings thus address an important question regarding higher antibody titers, especially against non-neutralizing non-canonical target antigens being potentially associated with worse outcomes in severe COVID-19 (Lee et al., 2020). The results suggest that higher antibody titers with particular isotypes directed against specific canonical as well as non-canonical antigens are associated with favorable outcomes in severe disease.

**Figure 2.**
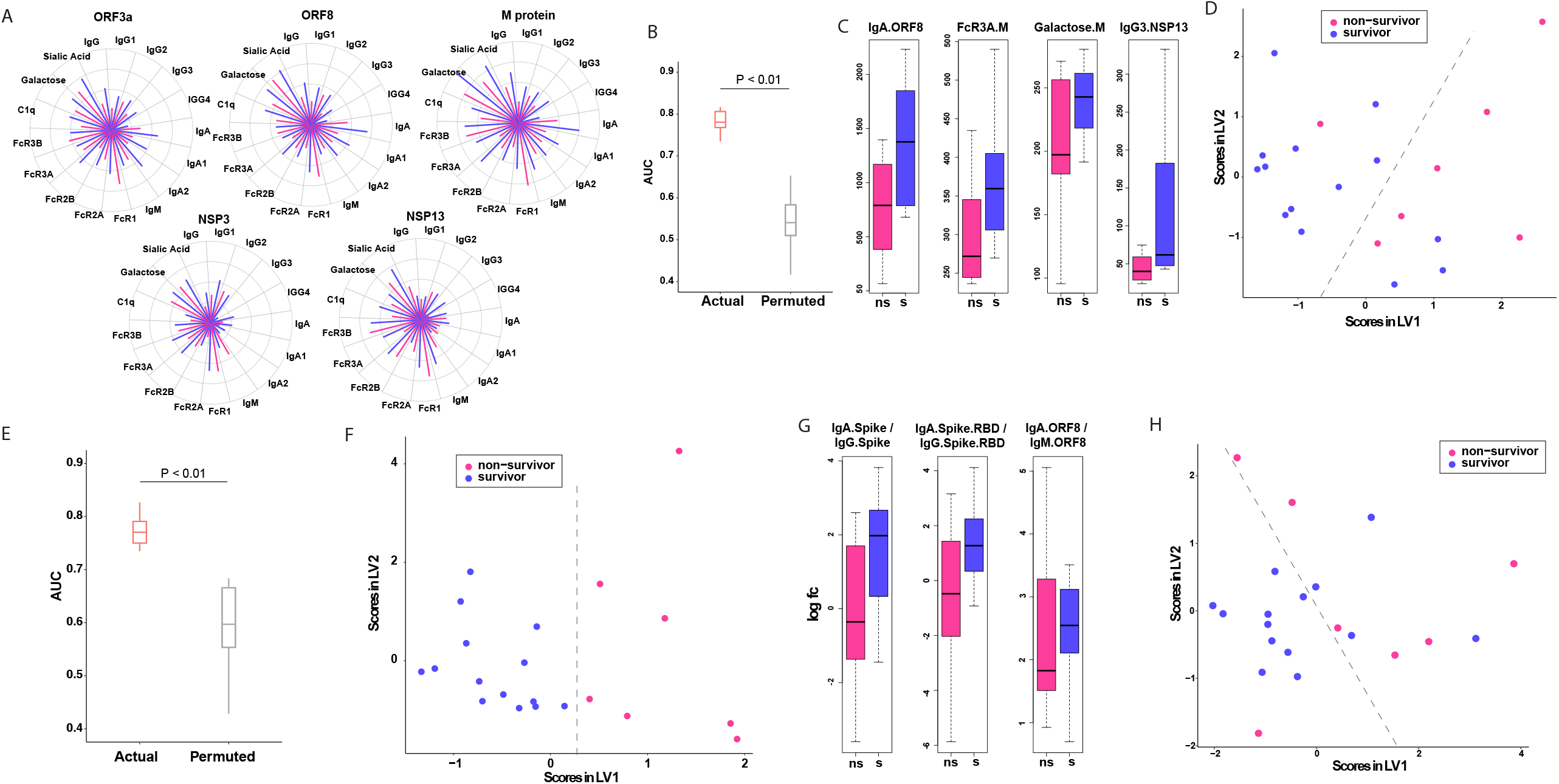
Multivariate antibody responses against non-canonical antigens independently predict severe COVID-19 outcomes. A. Polar plots illustrating measured antibody responses against non-canonical antigenic specificities orf3a, orf8, nsp3, nps13 and M. B. Performance of LASSO model to discriminate between survivors and non-survivors built using deep humoral profiles against non-canonical Ag specificities. Model performance is measured in a k-fold cross-validation framework with permutation testing. Actual denotes the performance of the model, built on real data. Permuted denotes performance of the model on shuffled data in a matched cross-validation framework (negative control). C. LASSO-selected features from model built using deep humoral profiles against non-canonical Ag specificities. D. PLS-DA using only the LASSO-selected features from the model in (C) to discriminate between survivors and non-survivors. E. Performance of LASSO model to discriminate between survivors and non-survivors built using deep humoral profiles against canonical and non-canonical Ag specificities. Model performance is measured in a k-fold cross-validation framework with permutation testing. Actual denotes the performance of the model, built on real data. Permuted denotes performance of the model on shuffled data in a matched cross-validation framework (negative control). F. LASSO-selected features from combined canonical and non-canonical antigenic specificities. G. Post-hoc feature selection based on the ratios of IgA to IgG and IgA to IgM. H. PLS-DA using only the ratios of IgA to IgG and IgA to IgM from the model in (G) to discriminate between survivors and non-survivors.

As noted above, IgA antibodies for both canonical (S-, S-RBD) and non-canonical (orf8) antigens were higher in survivors compared to non-survivors, (Figs. 1d, 2c). Further, increased galactosylation of both RBD- and M-specific Abs was associated with favorable outcomes (Figs. 1d, 2c). Thus, similar Ab profiles for both canonical and non-canonical specificities were associated with survival. This was corroborated, by constructing a predictive model by combining canonical and non-canonical specificities. As anticipated, the predictive performance of the model remained unchanged (Fig. 2e) and it highlighted a subset of the Ab features revealed by those based on canonical and non-canonical Ag specificities alone (Fig. 2f). Next, we examined whether the ratios of IgA/IgG or IgA/IgM antibodies directed against S, RBD and orf8 (antigens identified in the earlier analyses) were predictive of outcomes. This was a post-hoc analysis (Methods) where we focused on ratios of IgA/IgG antibodies for features identified in the canonical and non-canonical models (Fig. 2g). A multivariate PLS-DA visualization using just these 3 ratios discriminated between survivors and non-survivors (Figs 2h). These results reinforce the importance of IgA antibodies directed against both canonical and non-canonical specificities as important predictors of outcome in severe COVID-19. Overall, our results demonstrate that Abs with particular non-canonical antigen specificities are independently as informative as those Abs directed against the canonical S protein in predicting severe COVID-19 disease mortality outcomes.

### Antibodies directed against endemic CoV antigens as predictors of severe COVID-19 outcome

In the process of analyzing corresponding Ab profiles in pre-pandemic healthy controls we noticed significant levels of reactivities to specific non-canonical SARS-CoV-2 antigens, particularly nsp13 and nsp3 (Figs S1a-S1o and S2a-S2o). This was unlike Abs directed against canonical antigens including S which were, on the other hand, very close to or at baseline in these controls (Figs S1a-S1o and S2a-S2o). Given this surprising finding of Ab reactivity to nsp13 and nsp3 in sera of pre-pandemic individuals we hypothesized that these Abs may have been generated by prior infections of such individuals with eHCoVs and their cross-reactivity to SARS-Cov-2 nsp13 and nsp3 was a consequence of the greater conservation of these proteins across coronaviruses. To test this hypothesis, we analyzed the sequence similarity of nsp13 and S to corresponding antigens in SARS/MERS-CoV and eHCoVs (Fig. 3d), While S shares low sequence similarity to SARS-CoV and eHCoV S antigens, nsp13 has high sequence similarity to corresponding SARS/MERS-CoV and eHCoV antigens (Fig. 3d). Thus, antibodies directed against endemic eHCoV nsp13 may cross-react with their SARS-CoV2 homologs.

**Figure 3.**
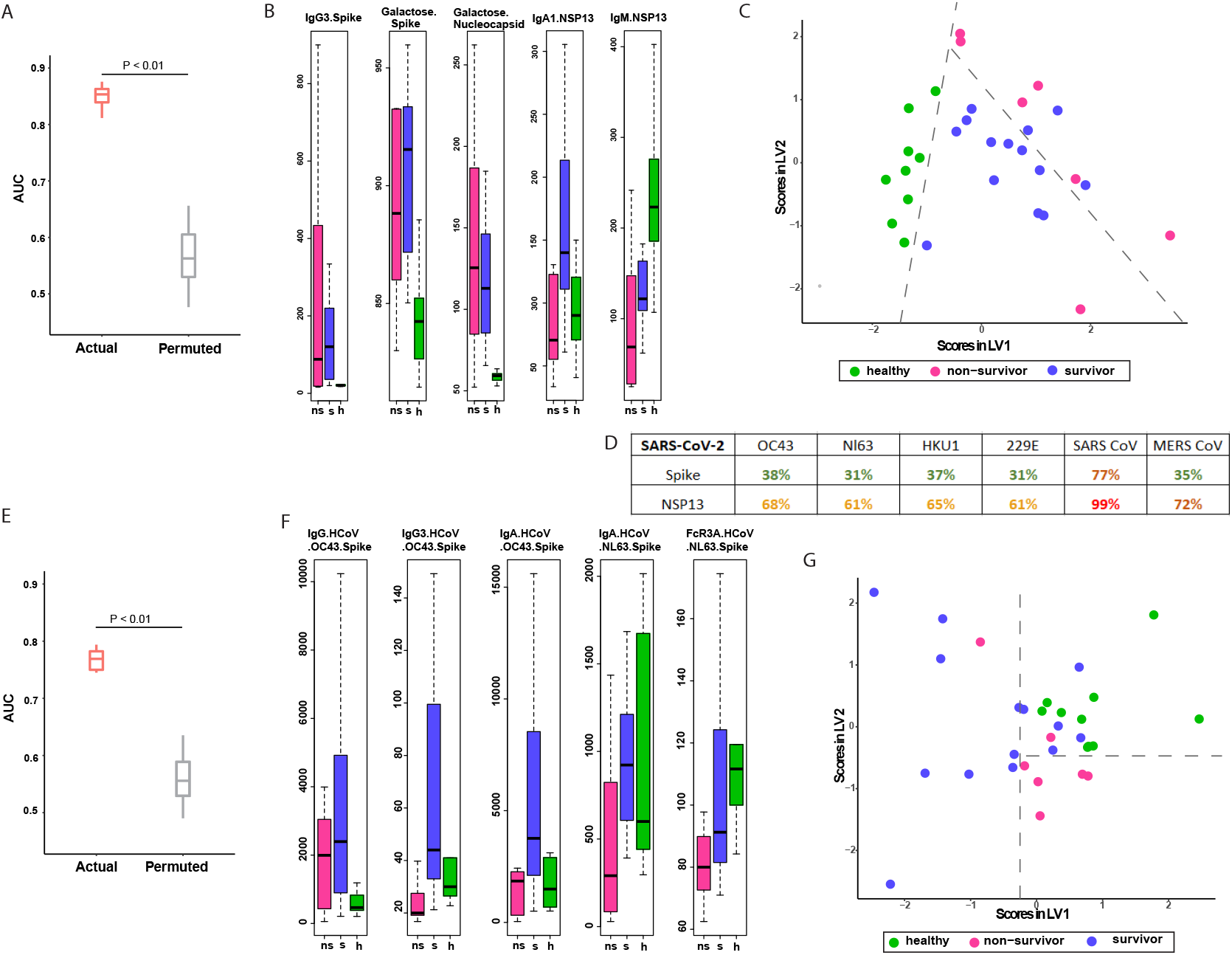
Antibodies directed against endemic CoV antigens as predictors of severe COVID-19 outcome. A. Performance of LASSO model to discriminate between survivors, non-survivors and healthy controls built using deep humoral profiles against canonical and non-canonical Ag specificities. Model performance (3-way) is measured in a k-fold cross-validation framework with permutation testing. Actual denotes the performance of the model, built on real data. Permuted denotes performance of the model on shuffled data in a matched cross-validation framework (negative control). B. LASSO-selected features from model built using deep humoral profiles against canonical and non-canonical Ag specificities. C. PLS-DA using only the LASSO-selected features from the model in (B) to discriminate between healthy controls, survivors, and non-survivors. D. Sequence similarities of SARS CoV2 Spike and NSP13 with corresponding homologs in OC43, NL63, HKU1, 229E, SARSCoV, and MERS CoV. E. Performance of LASSO model to discriminate between survivors, non-survivors and healthy controls built using deep humoral profiles against non-SARS-CoV2 Ag specificities. Model performance is measured in a k-fold cross-validation framework with permutation testing for outcome prediction between survivors, non-survivors, and healthy controls (3-way). Actual denotes the performance of the model, built on real data. Permuted denotes performance of the model on shuffled data in a matched cross-validation framework (negative control). F. LASSO-selected features from model built using deep humoral profiles against non-SARS-CoV2 Ag specificities. G. PLS-DA using only the LASSO-selected features from the model in (F) to discriminate between healthy controls, survivors, and non-survivors.

Given the possibility of pre-existing cross-reactive Abs to SARS-Cov-2 antigens we constructed a multivariate 3-way machine learning model that could discriminate not only between survivors and non-survivors, but also pre-pandemic healthy controls. Such a model might highlight additional Ab features pertaining to conserved non-canonical antigens that discriminate severe COVID-19 outcomes. The 3-way model performed even better than the previous 2-way models that discriminated between survivors and non-survivors (Fig. 3a). The LASSO model selected 5 features that reflected the intriguing trends described above. These features included anti-S IgG3, S-specific and N-specific antibody galactosylation as well as anti-nsp13 IgA1 and IgM (Fig. 3b). Analogous to our earlier analyses, a PLS-DA visualization also demonstrated that these 5 features were able to stratify survivors, non-survivors and healthy controls (Fig. 3c). Strikingly, while anti-S IgG3 was close to baseline in pre-pandemic healthy controls, the anti-nsp13 IgM was highest in pre-pandemic healthy controls and anti-nsp13 IgA1 was highest in survivors followed by healthy controls (Fig 3b). This raises the possibility that boosting of a pre-existing cross-reactive memory B cell response to nsp13 may account for the increased levels of these IgA Abs in severe COVID-19 patients and in turn their association with favorable outcomes.

To further test the possibility that boosting of pre-existing Abs to eHCoVs that may cross-react with SARS-CoV-2 antigens can associate with severe COVID-19 disease outcomes, we analyzed Abs directed against eHCoV antigens (OC43 S and NL63 S) and used that dataset to build a multivariate ML model. Remarkably, a multivariate model built only on Abs directed against OC43 S and NL63 S was significantly predictive of outcomes, and was also independently able to accurately discriminate between survivors, non-survivors and pre-pandemic healthy controls (Fig. 3e). Interestingly, while both survivors and non-survivors had significantly higher IgG antibodies to OC43 S than healthy controls, IgA, IgG3 antibodies and antibody binding to Fc receptor 3A were higher in survivors and healthy controls compared to non-survivors (Fig. 3f). IgA and IgG3 antibodies against OC43 S were also found to be higher in survivors relative to healthy controls. A PLS-DA visualization also demonstrated that these 5 markers were able to stratify survivors, non-survivors and healthy controls (Fig. 3g). These findings suggest that pre-existing Abs to canonical and non-canonical antigens of eHCoVs and their boosting, especially of particular isotypes and subclasses, by SARS-CoV-2 infection may have a beneficial role in favorable outcomes in severe COVID-19 disease.

## Discussion

Our study represents the deepest humoral profile of antibodies against canonical specificities and the first comprehensive profile of antibodies directed against non-canonical antigens as well as those directed against eHCoV antigens, in the context of severe COVID-19 disease. Critically, we found that multivariate models incorporating antibody responses against canonical antigens, and against non-canonical antigens were independently, equally discriminative of outcome. This demonstrates the importance of looking beyond the Spike and other canonical antigens (Zohar et al., 2020) and studying antibody profiles against these underexplored non-canonical specificities. Interestingly, similar molecular features of antibodies for both canonical and non-canonical specificities drove outcome bifurcation with survivors having more IgA and IgG3 isotypes, as well as higher antibody galactosylation. Importantly, these results demonstrate that within severe COVID-19 patients, outcome bifurcation can be accurately predicted at ICU admission based on IgA and IgG3 antibodies against canonical or non-canonical specificities. Notably, even with a small ICU cohort (n=21), we were able to obtain predictive models that were robust to cross-validation and permutation testing. This was made possible by deep high-dimensional Ab profiles corresponding with >100 Ab features/subject and the use of regularization-based machine learning to avoid overfitting (Ackerman et al., 2018; Das et al., 2020; Lu et al., 2020a; Sadanand et al., 2018; Suscovich et al., 2020).

Interestingly, pre-pandemic healthy controls were found to have humoral responses against SARS-CoV2 proteins with high sequence similarity with endemic coronaviruses, particularly nsp13. Notably, higher levels of IgA antibodies against nsp13 were found to be correlated with protection in severe COVID-19. Humoral profiles of eHCoV antibodies were also predictive of outcome bifurcation, with higher levels of IgA and IgG3 antibodies against OC43 and NL63 S being associated with survival.

Taken together, we demonstrate that antibodies targeting a range of specificities beyond just dominant and subdominant epitopes of the S protein, are predictive of outcome in severe COVID-19 disease. Further, in the context of a polyclonal response, examining the qualitative biophysical properties of these antibodies against non-canonical antigens i.e., not just IgG titers, but isotype/subclass composition, binding to Fc receptors and glycosylation profiles, which are all known to dictate Ab function, provides important discriminating features. While previous studies have focused solely on a small subset of SARS-CoV2 antigens, our study provides the first concrete evidence that that antibody responses against different subsets of antigens (canonical, non-canonical and endemic) are independently and equally discriminative of outcome. It has been speculated that higher antibody titers, especially against non-neutralizing targets, could be reflective of more severe disease and could also potentially be tied to ADE (Lee et al., 2020). Instead, our results suggest beneficial effects of IgA and IgG3 antibodies directed against canonical as well as non-canonical SARS-CoV-2 antigens. IgA antibodies function at mucosal surfaces and have previously been tied to vaccine-induced protection (Ackerman et al., 2018) in preventing death in severe COVID-19. Additionally, IgG3 is known to be a particularly potent inducer of antibody effector functions via higher affinity association with FcRs. Our findings suggest that class-switching to IgA and IgG3 isotypes directed not only at canonical but also non-canonical specificities may be important contributors to favorable outcomes in severe COVID-19. Further, complementary to earlier findings of aberrant glycosylation, specifically afucosylation, of S-specific Abs being correlated to disease severity (Larsen et al., 2021), we find here that differential glycosylation, specifically higher galactosylation, of Abs specific to both canonical and non-canonical antigens is associated with survival in severe COVID-19.

While there has been speculation regarding the roles of cross-reactive B and T cells as well as antibodies in driving outcome bifurcation for COVID-19 (Anderson et al., 2021; Le Bert et al., 2020; Shrock et al., 2020; (Loyal et al., 2021), our study identifies for the first time a range of multivariate humoral profiles of antibodies that can be robustly associated with outcome bifurcation in severe COVID-19. A recent study reported that not everyone exposed to SARS-CoV2 necessarily develops seropositivity, suggesting that some individuals clear sub-clinical infections (Swadling et al., 2021). In these individuals, pre-existing nsp13-specific T cells are expanded (Swadling et al., 2021). These findings strongly complement those described herein and collectively suggest that pre-existing nsp13-specific B cell responses are also likely boosted on exposure to SARS-CoV2 and in turn associate with favorable outcomes. Overall, our findings have major implications in the context of elucidating immunological states and associated intervention modalities in severe COVID-19. Our characterization of the protective nature of antibody responses against a broad panel of antigens has implications for the formulation of improved SARS-COV-2 as well as pan-coronavirus vaccines.

## Methods

### Sample preparation

Serum was obtained as described earlier and stored at −80C until used. Upon thawing before use, all samples were transferred to a 96 well U-bottom plate at an appropriate dilution for the intended probe (1:500 for IgG2, IgG3, IgG4, IgA, IgA1, IgA2, IgM, RCA I, SNA, C1Q & 1:2000 for IgG, IgG1, FcRs).

### Multiplexed Antigen-Specific Antibody Profiling

A multiplexed antigen-specific Ab profiling workflow was developed based on a protocol reported earlier (Brown et al., 2017). Briefly, pooled barcoded antigen-coupled beads were incubated with samples and then with fluorescently labelled probes. Following set of probes were used: 1) Antigen-specific subclass/isotype titers were measured using PE-labeled mouse-anti-human IgG1, IgG2, IgG3, IgG4, total IgG, IgA1, IgA2, IgM. 2) Antigen FcR/complement binding profiles were measured using biotinylated FcRs (FcR1, FcR2A, FcR2B, FcR3A, FcR3B) tetramerized with streptavidin-PE or PE-labeled complement (C1q). 3) Antigen-specific glycosylation profiles were measured using lectin-binding using PE-labeled lectins (SNA for sialic acid, RCA1 for galactosylation).

All antigens including SARS-CoV-2 Antigens (Spike [ImmuneTech IT-002-032p], Spike RBD [ImmuneTech IT-002-036p], Nucleocapsid [SinoBiological IT-002-036p], Orf3a [Bioworld NCP0026P], ORF8 [Bioworld NCP0025P], NSP3 [mybiosource MBS156024], NSP13 [mybiosource MBS2563852], M protein [mybiosource MBS156019]) and eHCoV antigens (HCoV-OC43 [SinoBiological 40607-V08B], HCoV-NL63 [SinoBiological 40604-V08B]) were coupled to Luminex MagPlex magnetic microsphere beads of different regions at a ratio of about 8ug of antigen per million beads using an EDC-NHS chemistry and then blocked with and stored until use in storage buffer (1XPBS, 0.1% BSA, 0.1% Tween) at 4C. PE coupled anti-Igs (Southern Biotech) were diluted from the stock vials to a concentration of 1ug/ml in 1XPBS. Biotinylated FcRs (Acro Biosystems) were reacted with Streptavidin-PE (Thermo Fisher) at a 4:1 molar ratio for 20 mins and then diluted to a concentration of 1ug/ml in 1XPBS. Rhodamine/Cy-3 coupled lectins (Vector Labs) were diluted in lectin buffer to a concentration of 20ug/ml.

Conjugated beads were diluted in assay buffer (1XPBS, 0.1% BSA) to make a working bead solution and added at a 1:9 sample:bead volume ratio in wells of a 96 well flat bottom plate, and incubated for 1 hour. Sample-bound beads were then washed twice in a wash buffer (PBS, 0.1% Tween) using a magnetic plate separator and resuspended in the appropriate probe buffer. Diluted probe solution was then added to the wells at a1:9 bead:probe volume ratio and incubated for 30 mins. All incubation steps were performed at room temperature on a plate shaker. Probe-bound beads were then washed twice with wash buffer and resuspended in Luminex MagPix drive fluid before reading on a Luminex MagPix instrument. All assays were performed in duplicate and a correlation coefficient of R^2^>0.8 was verified for technical replicability. An arithmetic mean of the two measured MFI values from the replicates is then used as the readout.

### Machine learning models to discriminate by outcome using humoral responses against different specificities (canonical, non-canonical and endemic CoV antigens)

Data was pre-processed to remove features with low values (mean MFI < 50). This was done in an unsupervised setting to avoid any biases. All features were centered and scaled (i.e., z-scored) to have a mean 0 and standard deviation 1.

We used a two-step machine learning model to identify a minimal set of predictive biomarkers of outcome. This comprised feature selection on the high-dimensional data (features >> number of subjects) using the least absolute shrinkage and selection operator (LASSO)(Tibshirani, 1996), followed by classification using the down-selected features using support vector machines(SVM)(Breiman, 2001). The use of LASSO (L1 regularization) helps prevent over-fitting on high dimensional data. This two-step procedure is similar to what has been successfully used earlier for high-dimensional humoral immune measurements(Ackerman et al., 2018; Lu et al., 2020b; Sadanand et al., 2018).

The performance of the models were evaluated in a rigorous 10-fold cross validation framework, and the significance of the models was quantified using permutation testing(Ojala and Garriga, 2010). The overall framework is analogous to what has been previously described(Ackerman et al., 2018; Lu et al., 2020b; Sadanand et al., 2018). Briefly, the dataset was split into 10 subsets – 9 subsets are used for training while the 10^th^ one is used for testing. Each subset served as the test set once, therefore each individual was in the test fold exactly once for each cross-validation run. For each test fold, LASSO-based feature selection was performed using the nine training folds. The coefficient for the LASSO penalty term (i.e., lambda for regularization) was determined via a second internal cross-validation using only the fold-specific training dataset. A fold-specific support vector machine (SVM) model was built using the LASSO-selected features and training data for that fold. This fold-specific classifier was subsequently used to predict the labels for the individuals in the test set for that fold. This process was repeated for each of the ten folds to generate a set of predicted outcomes for each individual. This was then compared to the true set of outcome labels to calculate a classification accuracy for that cross-validation replicate. We performed 100 independent ten-fold cross-validation replicates, to account for different ways in which the training and test folds can be split. This is a stringent and appropriate way of performing cross-validation, as both steps involved in the model (feature selection and subsequent classification using the selected features) are performed in a cross-validation setting with data held out. The significance of model performance was evaluated using permutation testing(Ojala and Garriga, 2010), by randomly shuffling the data with respect to the arm labels, within the cross-validation framework described above (i.e., a cross-validation framework matched to the actual model). To visualize the modules selected by the LASSO model on the whole dataset, we applied a partial least squares discriminant analysis (PLSDA). PLSDA is a supervised dimension reduction method, which transforms a new set of features that are linear combination of the original features and then fits a linear model via least squares using these new features. We carried out separate PLSDA analyses using down-selected features from applying LASSO on the whole dataset. PLSDA was applied for both 2 groups (between survivors and non-survivors) and 3 groups (among survivors, non-survivors, and healthy controls).

We analyzed the canonical, non-canonical and non-SARS-CoV2 specificities separately. We start with humoral responses for canonical (S/RBD/N) antigenic specificities to generate multivariate machine learning model for outcome (survivors vs non-survivors) bifurcation. We applied a similar approach on the non-canonical (orf3a/orf8/nsp3/nsp13/M) antigenic specificities for outcome bifurcation. We then combined both the canonical and non-canonical antigens and generate multivariate machine learning for outcome bifurcation between survivors and non-survivors. Then we also bring in the healthy controls along with the survivors and non-survivors and generate multivariate machine learning model for outcome prediction (3-way). We also built a multivariate machine learning model using endemic coronavirus antigenic specificities.

### Validation-cohort

To validate the robustness of biomarkers that we found in the Pittsburgh cohort, we only selected the canonical antigenic specificities that were common with the Boston cohort. Then we trained the machine learning model using only these common features of Pittsburgh cohort. The model (both feature selection and model fitting) was performed using only our cohort (Pittsburgh), and the model generation process was completely blinded to this second orthogonal validation cohort (Boston), ensuring that this is a true cross-prediction. Then we tested the performance of both the Pittsburgh and Boston cohort (common antigenic specificities) learned using the Pittsburgh cohort only. ROC curves for the LASSO selected features trained on both the cohort to compare and validate the performances.

### PLS-DA based visualization using IgA/IgG and IgA/IgM ratios

We performed a post-hoc greedy feature selection based on the ratios of IgA to IgG and IgA to IgM. We took the Spike, RBD, and Orf8 specificities for the ratios. We then performed a PLSDA on the selected ratios and to exhibit the discriminating power of these feature-ratios between survivors and non-survivors.

### Implementation of LASSO and PLS

LASSO was implemented using glmnet in R. If no feature was selected by LASSO in a specific fold for a given replicate, we randomly selected 5 features (only for that fold in that replicate) and use an ordinary least squares estimator. PLS was implemented using the plsr function in R.

## Figure Legends

**Supplementary Figure 1**

A-O. Dot plots showing deep humoral profiles against canonical (S/RBD/N) Ag specificities for survivors, non-survivors, and healthy controls (A-IgG, B-IgG1, C-IgG3, D-IgA, E-IgA1, F-IgA2, G-IgM, H-Galactose, I-Sialic acid, J-FcR2a, K-FcR3a, L-FcR3b, M-FcR1, N-FcR2b, O-C1Q.

P. Visualizing distribution of survivors and non-survivors for IgA.Spike and IgA2.Spike.RBD.

**Supplementary Figure 2**

A-O. Dot plots showing deep humoral profiles against non-canonical (orf3a/orf8/nsp3/nsp13/M) Ag specificities for survivors, non-survivors, and healthy controls (A-IgG, B-IgG1, C-IgG3, D-IgA, E-IgA1, F-IgA2, G-IgM, H-Galactose, I-Sialic acid, J-FcR2a, K-FcR3a, L-FcR3b, M-FcR1, N-FcR2b, O-C1Q.

P. LASSO-selected features from model built using deep humoral profiles against canonical and non-canonical Ag specificities.

**Supplementary Figure 3**

A-O. Dot plots showing deep humoral profiles against non-SARS-CoV2 (HCoVOC43, HCoVNL63) Ag specificities for survivors, non-survivors, and healthy controls (A-IgG, B-IgG1, C-IgG3, D-IgA, E-IgA1, F-IgA2, G-IgM, H-Galactose, I-Sialic acid, J-FcR2a, K-FcR3a, L-FcR3b, M-FcR1, N-FcR2b, O-C1Q.

## References

Ackerman, M.E., Das, J., Pittala, S., Broge, T., Linde, C., Suscovich, T.J., Brown, E.P., Bradley, T., Natarajan, H., Lin, S., et al. (2018). Route of immunization defines multiple mechanisms of vaccine-mediated protection against SIV. Nat Med 24, 1590–1598.

Atyeo, C., Fischinger, S., Zohar, T., Slein, M.D., Burke, J., Loos, C., McCulloch, D.J., Newman, K.L., Wolf, C., Yu, J., et al. (2020). Distinct Early Serological Signatures Track with SARS-CoV-2 Survival. Immunity 53, 524–532 e524.

Aydillo, T., Rombauts, A., Stadlbauer, D., Aslam, S., Abelenda-Alonso, G., Escalera, A., Amanat, F., Jiang, K., Krammer, F., Carratala, J., et al. (2021). Immunological imprinting of the antibody response in COVID-19 patients. Nat Commun 12, 3781.

Bain, W., Yang, H., Shah, F.A., Suber, T., Drohan, C., Al-Yousif, N., DeSensi, R.S., Bensen, N., Schaefer, C., Rosborough, B.R., et al. (2021). COVID-19 versus Non-COVID-19 Acute Respiratory Distress Syndrome: Comparison of Demographics, Physiologic Parameters, Inflammatory Biomarkers, and Clinical Outcomes. Ann Am Thorac Soc 18, 1202–1210.

Bartsch, Y.C., Fischinger, S., Siddiqui, S.M., Chen, Z., Yu, J., Gebre, M., Atyeo, C., Gorman, M.J., Zhu, A.L., Kang, J., et al. (2021). Discrete SARS-CoV-2 antibody titers track with functional humoral stability. Nat Commun 12, 1018.

Breiman, L. (2001). Random Forests. Machine Learning 45, 5–32.

Brown, E.P., Dowell, K.G., Boesch, A.W., Normandin, E., Mahan, A.E., Chu, T., Barouch, D.H., Bailey-Kellogg, C., Alter, G., and Ackerman, M.E. (2017). Multiplexed Fc array for evaluation of antigen-specific antibody effector profiles. J Immunol Methods 443, 33–44.

Bye, A.P., Hoepel, W., Mitchell, J.L., Jegouic, S., Loureiro, S., Sage, T., Vidarsson, G., Nouta, J., Wuhrer, M., de Taeye, S., et al. (2021). Aberrant glycosylation of anti-SARS-CoV-2 spike IgG is a prothrombotic stimulus for platelets. Blood 138, 1481–1489.

Carvalho, T., Krammer, F., and Iwasaki, A. (2021). The first 12 months of COVID-19: a timeline of immunological insights. Nat Rev Immunol 21, 245–256.

Chakraborty, S., Gonzalez, J., Edwards, K., Mallajosyula, V., Buzzanco, A.S., Sherwood, R., Buffone, C., Kathale, N., Providenza, S., Xie, M.M., et al. (2020). Proinflammatory IgG Fc structures in patients with severe COVID-19. medRxiv.

Cillo, A.R., Somasundaram, A., Shan, F., Cardello, C., Workman, C.J., Kitsios, G.D., Ruffin, A., Kunning, S., Lampenfeld, C., Onkar, S., et al. (2021). Bifurcated monocyte states are predictive of mortality in severe COVID-19. bioRxiv.

Das, J., Devadhasan, A., Linde, C., Broge, T., Sassic, J., Mangano, M., O’Keefe, S., Suscovich, T., Streeck, H., Irrinki, A., et al. (2020). Mining for humoral correlates of HIV control and latent reservoir size. PLoS Pathog 16, e1008868.

Gordon, D.E., Jang, G.M., Bouhaddou, M., Xu, J., Obernier, K., White, K.M., O’Meara, M.J., Rezelj, V.V., Guo, J.Z., Swaney, D.L., et al. (2020). A SARS-CoV-2 protein interaction map reveals targets for drug repurposing. Nature 583, 459–468.

Grifoni, A., Weiskopf, D., Ramirez, S.I., Mateus, J., Dan, J.M., Moderbacher, C.R., Rawlings, S.A., Sutherland, A., Premkumar, L., Jadi, R.S., et al. (2020). Targets of T Cell Responses to SARS-CoV-2 Coronavirus in Humans with COVID-19 Disease and Unexposed Individuals. Cell 181, 1489–1501 e1415.

Guo, L., Wang, Y., Kang, L., Hu, Y., Wang, L., Zhong, J., Chen, H., Ren, L., Gu, X., Wang, G., et al. (2021). Cross-reactive antibody against human coronavirus OC43 spike protein correlates with disease severity in COVID-19 patients: a retrospective study. Emerg Microbes Infect 10, 664–676.

Hachim, A., Kavian, N., Cohen, C.A., Chin, A.W.H., Chu, D.K.W., Mok, C.K.P., Tsang, O.T.Y., Yeung, Y.C., Perera, R., Poon, L.L.M., et al. (2020). ORF8 and ORF3b antibodies are accurate serological markers of early and late SARS-CoV-2 infection. Nat Immunol 21, 1293–1301.

Hicks, J., Klumpp-Thomas, C., Kalish, H., Shunmugavel, A., Mehalko, J., Denson, J.P., Snead, K.R., Drew, M., Corbett, K.S., Graham, B.S., et al. (2021). Serologic Cross-Reactivity of SARS-CoV-2 with Endemic and Seasonal Betacoronaviruses. J Clin Immunol 41, 906–913.

Hoepel, W., Chen, H.J., Geyer, C.E., Allahverdiyeva, S., Manz, X.D., de Taeye, S.W., Aman, J., Mes, L., Steenhuis, M., Griffith, G.R., et al. (2021). High titers and low fucosylation of early human anti-SARS-CoV-2 IgG promote inflammation by alveolar macrophages. Sci Transl Med 13.

Iwasaki, A., and Yang, Y. (2020). The potential danger of suboptimal antibody responses in COVID-19. Nat Rev Immunol 20, 339–341.

Khoury, D.S., Cromer, D., Reynaldi, A., Schlub, T.E., Wheatley, A.K., Juno, J.A., Subbarao, K., Kent, S.J., Triccas, J.A., and Davenport, M.P. (2021). Neutralizing antibody levels are highly predictive of immune protection from symptomatic SARS-CoV-2 infection. Nat Med 27, 1205–1211.

Lamere, M.W., Moquin, A., Lee, F.E., Misra, R.S., Blair, P.J., Haynes, L., Randall, T.D., Lund, F.E., and Kaminski, D.A. (2011). Regulation of antinucleoprotein IgG by systemic vaccination and its effect on influenza virus clearance. J Virol 85, 5027–5035.

Larsen, M.D., de Graaf, E.L., Sonneveld, M.E., Plomp, H.R., Nouta, J., Hoepel, W., Chen, H.J., Linty, F., Visser, R., Brinkhaus, M., et al. (2021). Afucosylated IgG characterizes enveloped viral responses and correlates with COVID-19 severity. Science 371.

Lee, W.S., Wheatley, A.K., Kent, S.J., and DeKosky, B.J. (2020). Antibody-dependent enhancement and SARS-CoV-2 vaccines and therapies. Nat Microbiol 5, 1185–1191.

Loyal, L., Braun, J., Henze, L., Kruse, B., Dingeldey, M., Reimer, U., Kern, F., Schwarz, T., Mangold, M., Unger, C., et al. (2021). Cross-reactive CD4(+) T cells enhance SARS-CoV-2 immune responses upon infection and vaccination. Science 374, eabh1823.

Lu, L.L., Das, J., Grace, P.S., Fortune, S.M., Restrepo, B.I., and Alter, G. (2020a). Antibody Fc Glycosylation Discriminates Between Latent and Active Tuberculosis. J Infect Dis.

Lu, L.L., Das, J., Grace, P.S., Fortune, S.M., Restrepo, B.I., and Alter, G. (2020b). Antibody Fc Glycosylation Discriminates Between Latent and Active Tuberculosis. J Infect Dis, In press.

Ng, W.H., Tipih, T., Makoah, N.A., Vermeulen, J.G., Goedhals, D., Sempa, J.B., Burt, F.J., Taylor, A., and Mahalingam, S. (2021). Comorbidities in SARS-CoV-2 Patients: a Systematic Review and Meta-Analysis. mBio 12.

Ojala, M., and Garriga, G.C. (2010). Permutation Tests for Studying Classifier Performance. J Mach Learn Res 11, 1833–1863.

Rodebaugh, T.L., Frumkin, M.R., Reiersen, A.M., Lenze, E.J., Avidan, M.S., Miller, J.P., Piccirillo, J.F., Zorumski, C.F., and Mattar, C. (2021). Acute Symptoms of Mild to Moderate COVID-19 Are Highly Heterogeneous Across Individuals and Over Time. Open Forum Infect Dis 8, ofab090.

Sadanand, S., Das, J., Chung, A.W., Schoen, M.K., Lane, S., Suscovich, T.J., Streeck, H., Smith, D.M., Little, S.J., Lauffenburger, D.A., et al. (2018). Temporal variation in HIV-specific IgG subclass antibodies during acute infection differentiates spontaneous controllers from chronic progressors. AIDS 32, 443–450.

Sadarangani, M., Marchant, A., and Kollmann, T.R. (2021). Immunological mechanisms of vaccine-induced protection against COVID-19 in humans. Nat Rev Immunol 21, 475–484.

Shrock, E., Fujimura, E., Kula, T., Timms, R.T., Lee, I.H., Leng, Y., Robinson, M.L., Sie, B.M., Li, M.Z., Chen, Y., et al. (2020). Viral epitope profiling of COVID-19 patients reveals cross-reactivity and correlates of severity. Science 370.

Suscovich, T.J., Fallon, J.K., Das, J., Demas, A.R., Crain, J., Linde, C.H., Michell, A., Natarajan, H., Arevalo, C., Broge, T., et al. (2020). Mapping functional humoral correlates of protection against malaria challenge following RTS,S/AS01 vaccination. Sci Transl Med 12.

Swadling, L., Diniz, M.O., Schmidt, N.M., Amin, O.E., Chandran, A., Shaw, E., Pade, C., Gibbons, J.M., Le Bert, N., Tan, A.T., et al. (2021). Pre-existing polymerase-specific T cells expand in abortive seronegative SARS-CoV-2. Nature.

Tibshirani, R. (1996). Regression shrinkage and selection via the lasso. Journal of the Royal Statistical Society: Series B (Methodological) 58, 267–288.

To, K.K., Zhang, A.J., Hung, I.F., Xu, T., Ip, W.C., Wong, R.T., Ng, J.C., Chan, J.F., Chan, K.H., and Yuen, K.Y. (2012). High titer and avidity of nonneutralizing antibodies against influenza vaccine antigen are associated with severe influenza. Clin Vaccine Immunol 19, 1012–1018.

Zohar, T., Loos, C., Fischinger, S., Atyeo, C., Wang, C., Slein, M.D., Burke, J., Yu, J., Feldman, J., Hauser, B.M., et al. (2020). Compromised Humoral Functional Evolution Tracks with SARS-CoV-2 Mortality. Cell 183, 1508–1519 e1512.

