## Supplementary figures and images for "Antibodies targeting conserved non-canonical antigens and endemic coronaviruses associate with favorable outcomes in severe COVID-19"

### Supplementary Figures

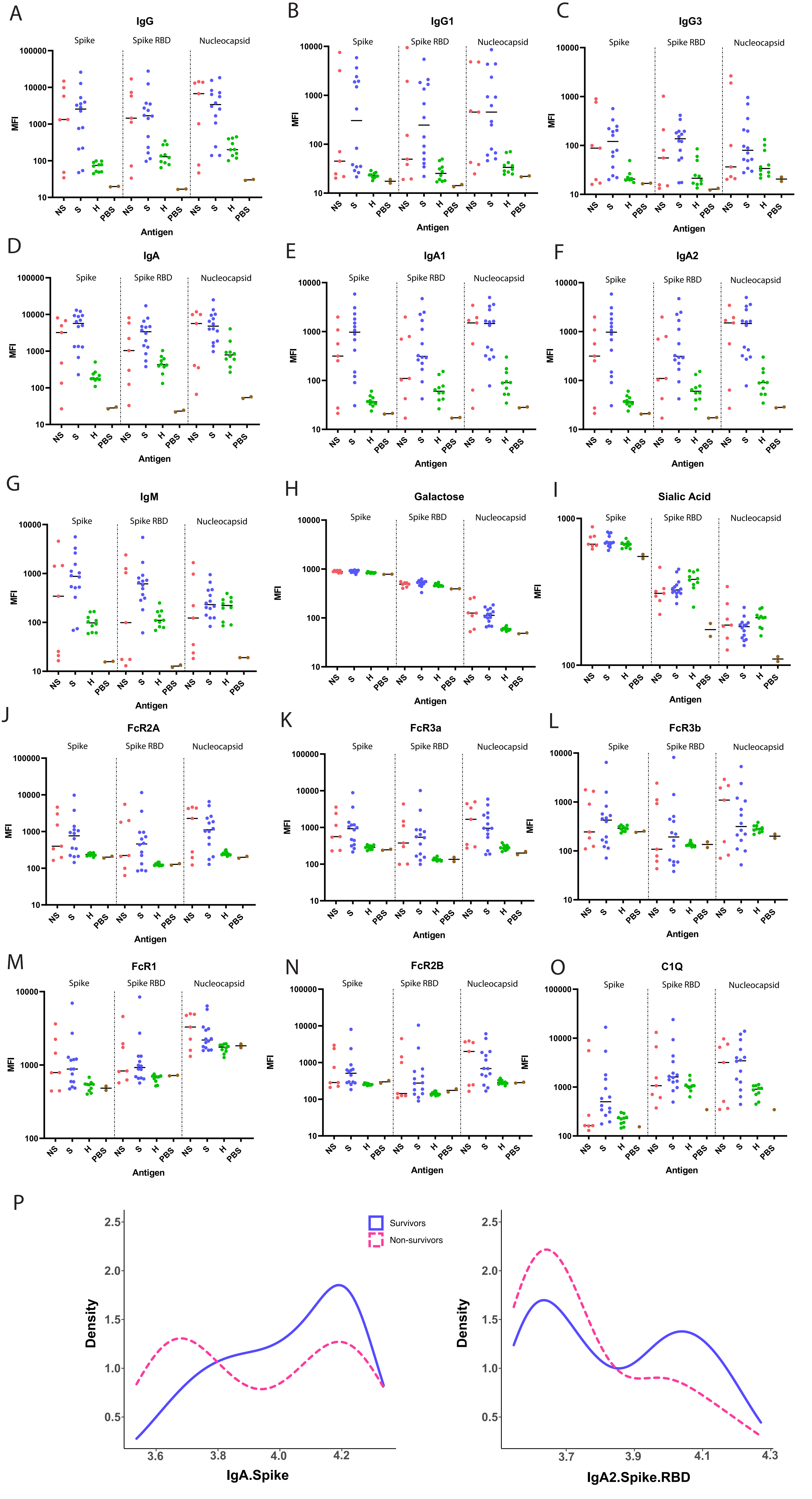

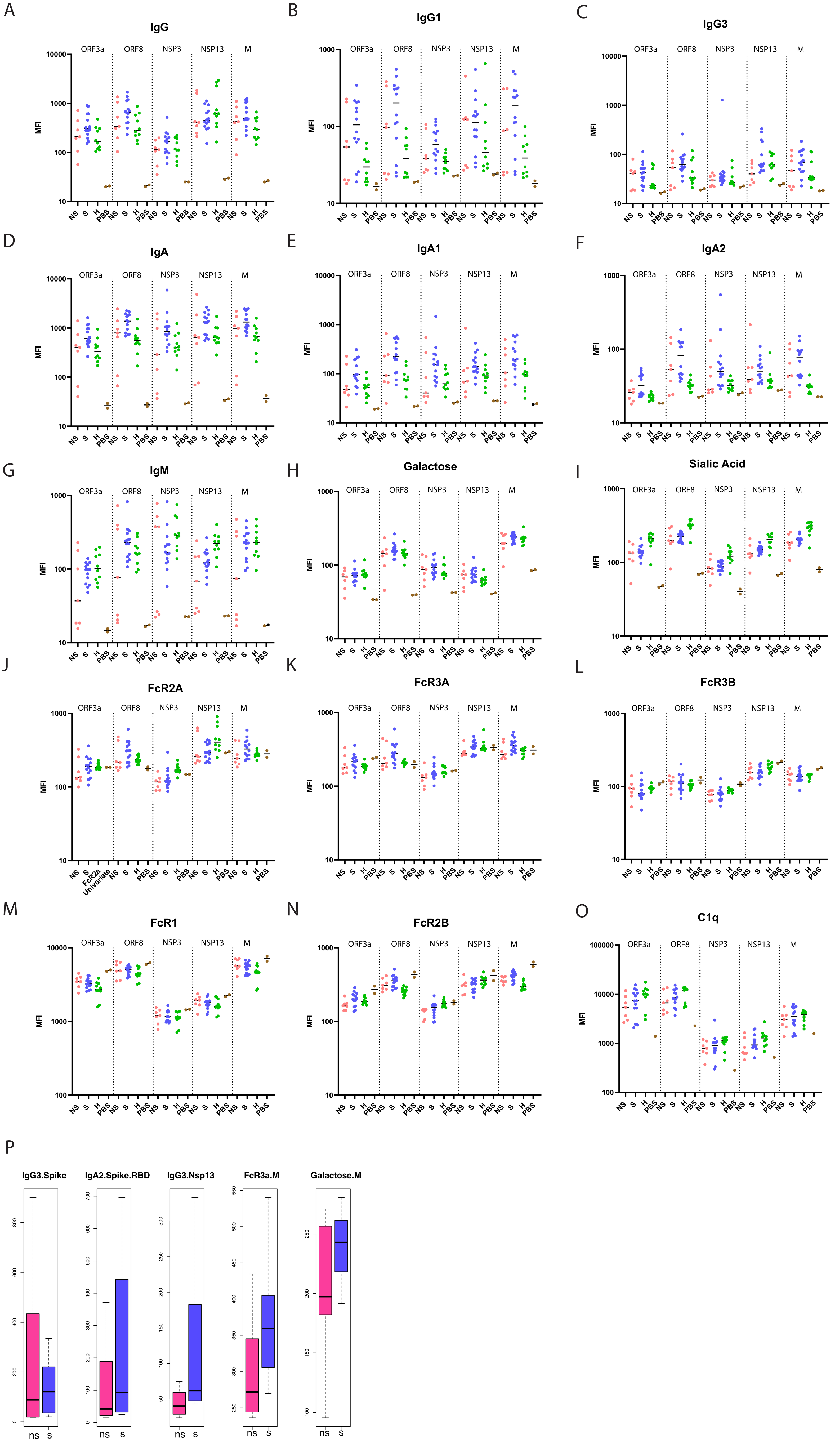

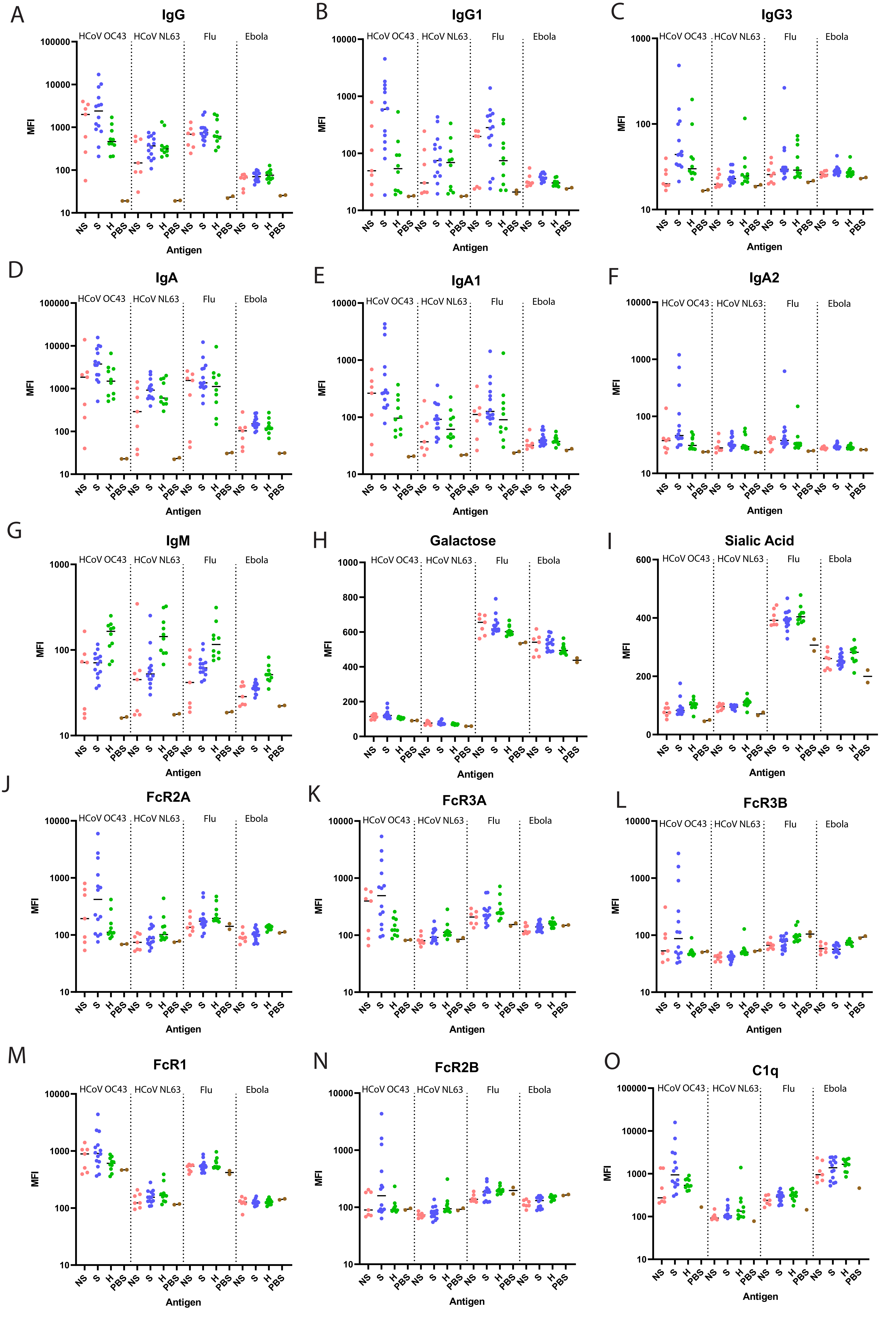
